# Network-targeted, multi-site direct cortical stimulation enhances working memory by modulating phase lag of low frequency oscillations

**DOI:** 10.1101/514554

**Authors:** Sankaraleengam Alagapan, Justin Riddle, Wei Angel Huang, Eldad Hadar, Hae Won Shin, Flavio Fröhlich

## Abstract

Working memory, an important component of cognitive control, is supported by the coordinated activation of a network of cortical regions in the frontal and parietal cortices. Oscillations in theta and alpha frequency bands are thought to coordinate these network interactions. Thus, targeting multiple nodes of the network with brain stimulation at the frequency of interaction may be an effective means of modulating working memory. We tested this hypothesis by identifying regions that are functionally connected in theta and alpha frequency bands and intracranially stimulating both regions simultaneously in participants undergoing invasive monitoring. We found that in-phase stimulation resulted in improvement in performance compared to sham stimulation. In contrast, anti-phase stimulation did not affect performance. In-phase stimulation resulted in decreased phase lag between regions within working memory network while anti-phase stimulation resulted in increased phase lag suggesting that shorter phase lag in oscillatory connectivity may lead to better performance. The results support the idea that phase lag may play a key role in information transmission across brain regions. More broadly, brain stimulation strategies that aim to improve cognition may be better served targeting multiple nodes of brain networks.

## Introduction

Working memory (WM) is an important component of cognition and supports higher cognitive functions in humans like fluid intelligence, decision making and learning. Impairment of WM is observed in many psychiatric and neurological disorders [1–3] and is often not addressed by current treatment strategies. Thus, approaches that can improve WM are required. The neural substrates of WM are spread across frontal, cingulate and parietal cortices [4–7] and are thought to be coordinated by cortical oscillations. Theta (4 – 8 Hz) and alpha (8 – 12 Hz) oscillations are known to play a critical role in WM [8–11]. Given the spatially distributed nature of processing that takes place during WM tasks, the interaction between different regions that underlies WM can be captured in functional and effective connectivity analyses. Neuroimaging studies have revealed that fronto-parietal connectivity is a key functional component of WM in the brain [12–14] and some studies have found connectivity between frontal and temporal regions to be correlated with WM task performance [15, 16]. Electroencephalography (EEG) and magnetoencephalography (MEG) studies have shown that fronto-parietal connectivity may be characterized by interactions in different oscillatory frequency bands. Alpha band phase synchronization in fronto-parietal regions has been shown to be modulated by WM load [17, 18]. Theta band connectivity has been shown to increase with increased central executive demands [19, 20]. Deficits in WM are common in many neurological and psychiatric disorders in which connectivity is also altered [21–24]. Taken together, the neural substrate for WM is a network of brain regions and thus, any strategy that targets WM may be better-served by engaging multiple nodes of the network.

Noninvasive brain stimulation methods like transcranial magnetic stimulation (TMS) [25–28], transcranial direct current stimulation (tDCS) [29–31] and transcranial alternating current stimulation (tACS) [32–36] have allowed causal perturbations of specific regions or activity signatures involved in WM. More specifically, rhythmic TMS (rTMS), in which a periodic pulse train is applied, and tACS, in which a continuous sinusoidal alternating current is applied, allow for targeting neural oscillations by matching the stimulation frequency to the frequency of oscillations [37]. RTMS has been shown to improve WM performance when applied at theta frequency [38–40]. TACS in theta frequency band also leads to improvements in WM performance [33, 36]. Most of these studies have focused on stimulating a single region. In contrast, studies in which multiple regions of WM network are targeted have yielded important insights into functional network properties. TACS studies have shown that stimulating fronto-parietal network using waveforms that have 0° phase offset (in-phase stimulation) result in improvement of WM performance while stimulating networks using waveforms that have 180° phase offset (anti-phase stimulation) result in deterioration of performance [34, 35]. In-phase stimulation was hypothesized to cause synchronization of the fronto-parietal networks while anti-phase stimulation was is hypothesized to cause de-synchronization. Neuroimaging during stimulation indicated increased blood oxygenation level dependent (BOLD) signal in WM regions during in-phase stimulation while functional connectivity increased with both in-phase stimulation and anti-phase stimulation [35]. The BOLD signal does not have milli-second temporal resolution and thus precluded any analysis of the changes in oscillatory network activity.

Compared to transcranial electric stimulation, direct cortical stimulation (DCS), in which electrical stimulation is applied directly on the cortical surface, offers higher spatial specificity. Additionally, intracranial EEG (iEEG) provides higher spatial resolution relative to EEG or MEG as well as higher temporal resolution relative to functional neuroimaging. Thus, by combining DCS and iEEG, it is possible to dissect functional networks with high spatio-temporal precision. This approach has been used for causally perturbing the electrophysiological and anatomical substrates of episodic memory [41–43], memory consolidation [44], and face processing [45, 46]. DCS has also been used to target networks engaged in spatial memory, albeit stimulation resulted in impairment of performance [47]. In another study, direct stimulation of bilateral hippocampal regions with in-phase and anti-phase stimulation resulted in trend-level changes in performance [48]. Using this approach, we have shown that frequency-matched DCS of a region (left superior frontal gyrus) that exhibited low frequency oscillatory activity results in working memory improvement [49]. Here, we extended our stimulation protocol to target networks underlying working memory by stimulating two functionally connected regions simultaneously. We used a measure of phase synchronization, the weighted phase lag index, to identify regions that are functionally connected in alpha and theta frequency bands during a Sternberg WM task. We applied periodic pulse stimulation in-phase and anti-phase, matched to the frequency of functional interactions, to the two functionally connected regions, and compared the performance against sham stimulation. We hypothesized that in-phase stimulation would result in an increase in oscillatory functional connectivity relative to sham and thereby improve WM performance while anti-phase stimulation would result in a decrease in oscillatory functional connectivity relative to sham and thereby impair WM performance. While in-phase stimulation improved performance, anti-phase stimulation did not impair performance relative to sham. Analysis of functional connectivity properties in atlas-based WM (aWM) network revealed that functional connectivity was increased by both in-phase and anti-phase stimulation. However, in-phase stimulation decreased phase lag relative to sham between regions within the aWM network while anti-phase stimulation increased phase lag relative to sham suggesting a non-linear relationship between the phase lag of connections within a network and performance.

## Results

We performed network-targeted stimulation (Figure 1A) in 3 participants implanted with subdural strips and stereo EEG electrodes for epilepsy surgery planning. The electrodes covered bilateral frontal, parietal and temporal cortices (Figure S1). Participants performed a Sternberg working memory task (Figure 1B) in a baseline session and a stimulation session. In the stimulation session, trains of biphasic pulses were applied to two pairs of electrodes. Stimulation was applied in-phase, in which stimulation was applied simultaneously between the two electrode pairs, and anti-phase, in which stimulation applied between one pair of electrodes was temporally offset from the other electrode pair by half the inter-pulse-interval of the pulse train (Figure 1C). Stimulation was applied during the encoding epoch of the Sternberg task. Sham stimulation, in which no DCS was applied, was used as a control as iEEG participants are unable to tell when stimulation is applied. The three stimulation conditions (in-phase, anti-phase, and sham) were randomly interleaved for each task block.

**Figure 1:**
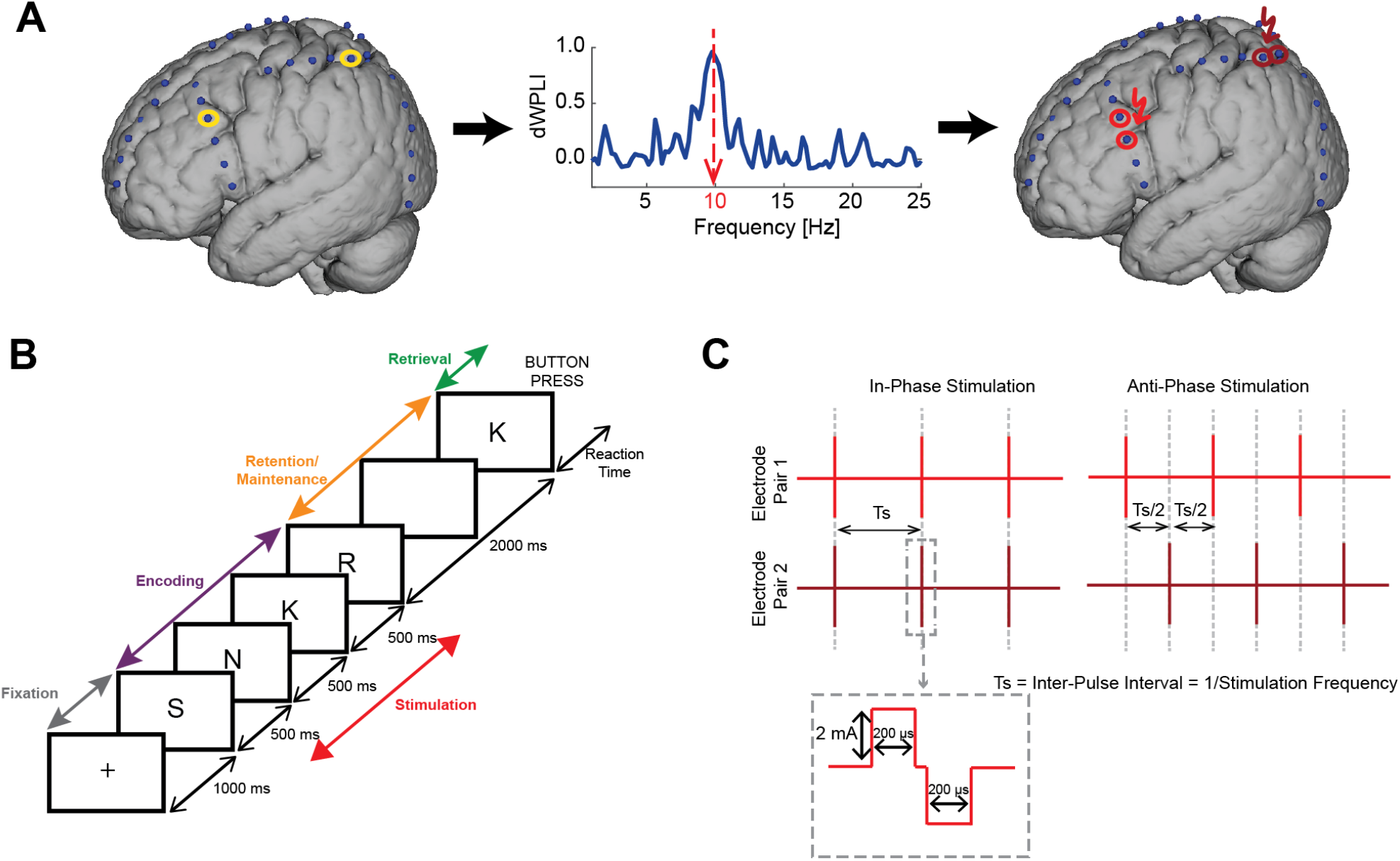
Schematic of Network-Targeted Stimulation. A. Intracranial EEG data from implanted electrodes, collected when participants performed WM task, are processed to identify functionally connected regions that are then targeted with direct cortical stimulation. B. Sternberg working memory task depicting the different epochs and timing of components of each epoch C. The stimulation paradigms used in the study. Each vertical red line denotes a biphasic pulse. In-phase stimulation consists of pulses applied simultaneously to functionally connected regions without any phase offset (time delay). Anti-phase stimulation consists of pulses applied with a phase offset of 180° (time delay of half the inter-stimulus interval Ts). Dotted lines are provided for visual guidance

In the baseline session, the WM load, defined as the number of items to be held in WM, was varied pseudo randomly for each trial. The WM load for each participant was titrated according to performance in a short practice session (3, 5 for P1; 5, 7 for P2 and P3). Chi-squared test did not reveal any significant influence of list length on accuracy (χ^2^ = 0.434, df = 2, p = 0.805). Analysis of reaction time did not reveal any significant influence of list length (Linear mixed effects model with list length as fixed factor and participant as random factor; F_2, 175.51_ = 0.630, p = 0.534). The reaction time and accuracy for individual participants are shown in Figure S2.

Analysis of functional connectivity using debiased weighted phase lag index (dWPLI) revealed oscillatory interactions in theta and alpha frequency bands. DWPLI measures the degree of consistency of phase lag between two signals and is not affected by volume conduction [50] making it an effective tool for identifying functional interactions in iEEG. In P1, electrodes that exhibited connectivity within the left frontal regions (superior frontal gyrus and inferior precentral gyrus) in theta band (4 Hz) were chosen. In P2, electrodes in the left frontal and parietal regions (inferior frontal junction and superior parietal lobule) that exhibited interactions in alpha band interactions were chosen. In P3, no strong functional interactions were observed (apart from the interactions between neighboring electrodes). Therefore, we chose electrodes that were in the putative WM network in the right hemisphere (middle frontal gyrus and superior intraparietal sulcus). We chose 10 Hz as stimulation frequency for P3 as alpha band synchronization between frontal and parietal regions has been shown to impact WM [17, 18]. The mean dWPLI for the electrodes chosen are shown in Figure 2A. Post-hoc analysis of spatial proximity of the chosen stimulation electrodes to canonical WM network identified from a meta-analytic atlas [51] revealed that both the electrode pairs in P2 and P3 were in or near regions active during WM (Figure 2B). In P1, one electrode pair was near the inferior frontal junction, another prominent WM region [7] and one pair was on the superior frontal gyrus, a region that we have previously demonstrated to be involved in WM [49].

**Figure 2:**
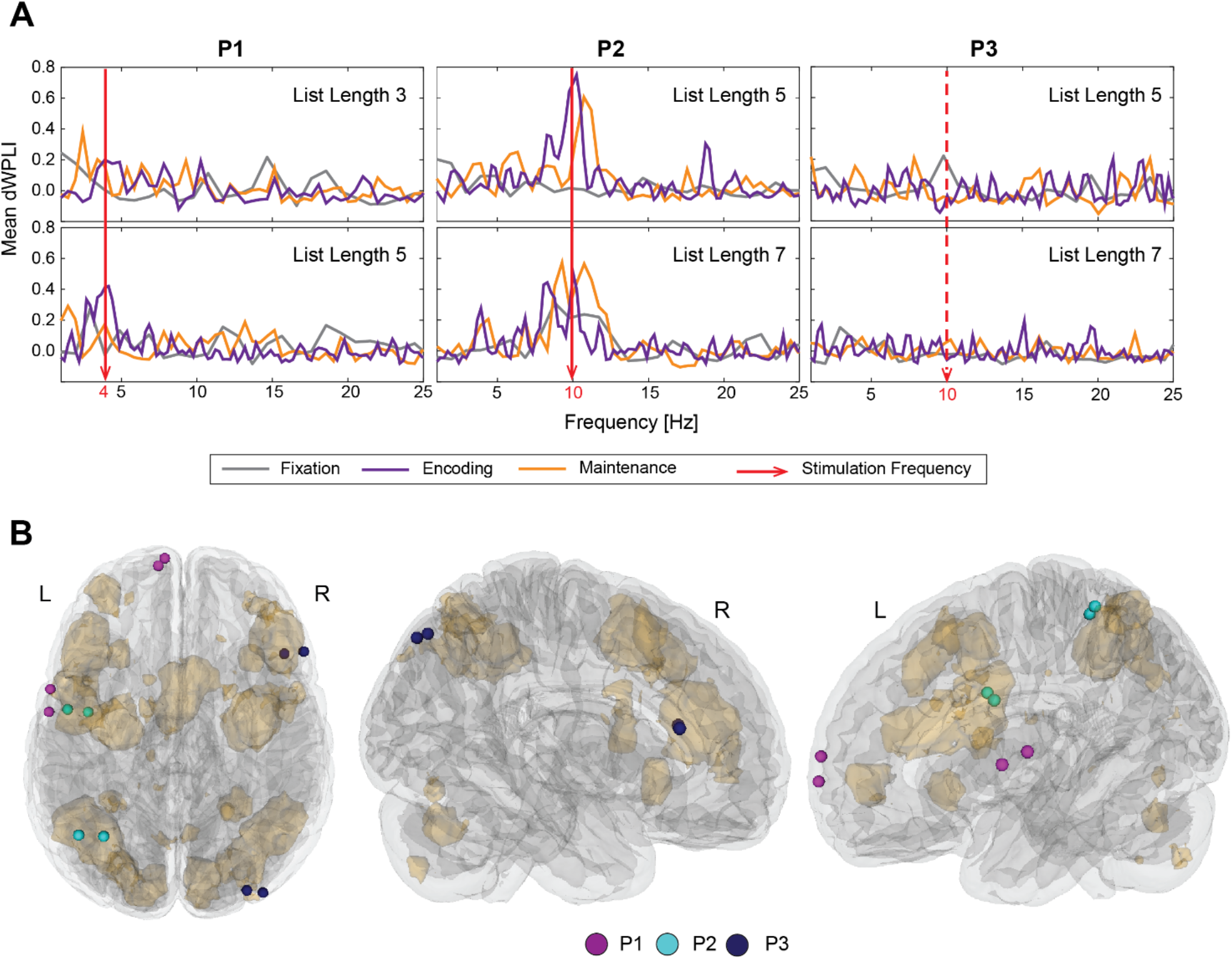
Functional connectivity of stimulation electrodes. A. Mean dWPLI for the stimulation electrodes for the different cognitive loads and the different epochs. B. The anatomical locations of the identified stimulation electrodes for the three participants. The beige shaded regions denote WM regions identified from meta-analyses of functional neuroimaging studies.

In stimulation session, participants performed the Sternberg task again but with only one level of WM load. Stimulation was applied between pairs of electrodes identified in the baseline session during the encoding epoch. In-phase stimulation resulted in increased accuracy relative to sham in all 3 participants (Figure 3, Top). Chi-squared test with all three conditions revealed a statistically significant association between condition and trial accuracy (χ^2^ = 7.315, df = 2, p = 0.026). Further pairwise comparisons revealed that inphase stimulation increased accuracy relative to sham (χ^2^ = 6.429 df = 1, p = 0.011), but there was no difference in accuracy for anti-phase relative to sham (χ^2^ = 1.0913 df = 1, p = 0.296) or in-phase relative to anti-phase (χ^2^ = 1.847 df = 1, p = 0.174). Thus, network-targeted stimulation improved WM accuracy but only when both electrode pairs were stimulated simultaneously without phase-lag. Analysis of reaction time did not reveal any statistically significant effect of stimulation condition (Figure 3, Bottom, Linear mixed model with fixed factor stimulation condition and random factor participant; F_2,223.42_ = 0.545, p = 0.581).

**Figure 3:**
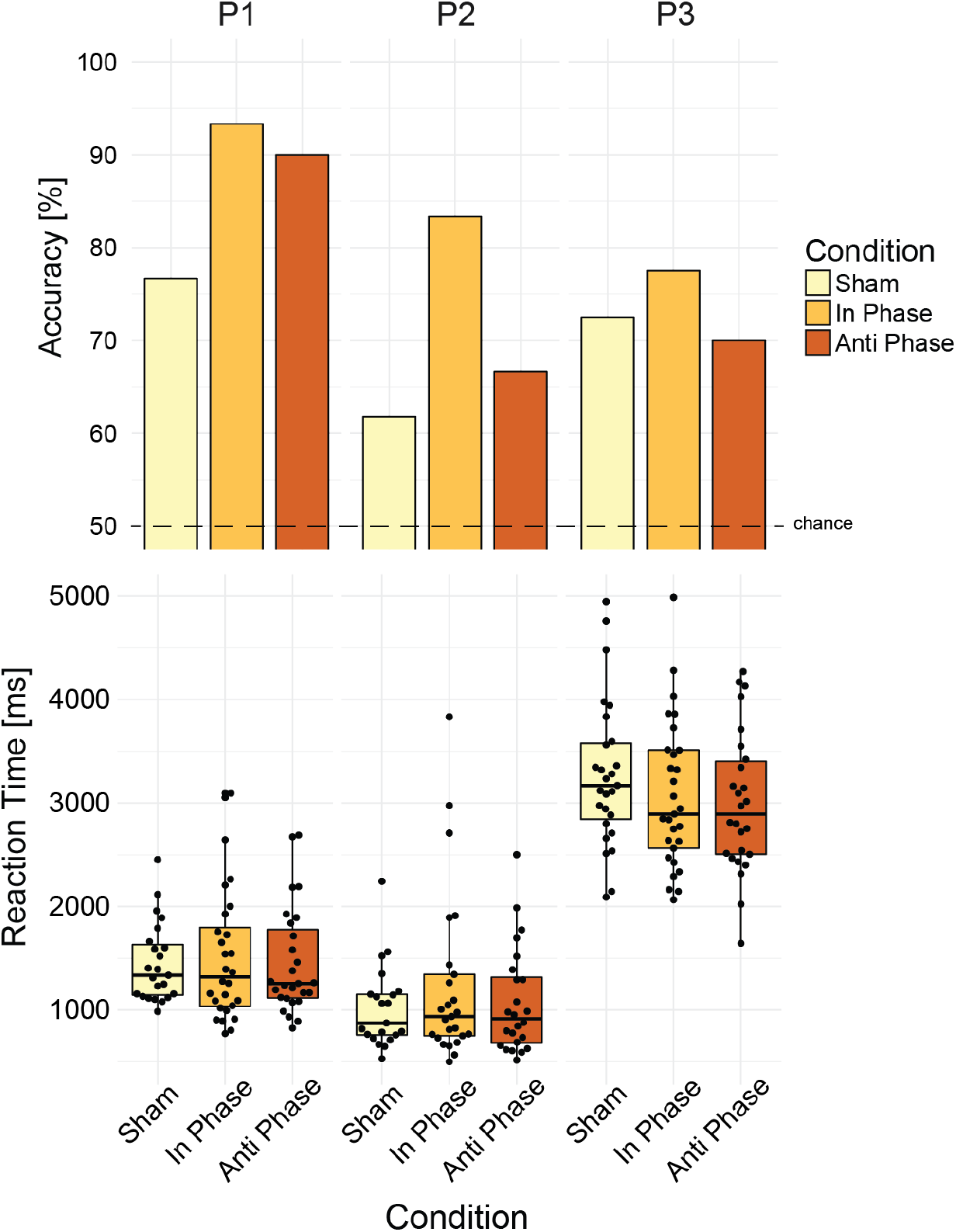
Effect of network-targeted stimulation on WM performance. In-phase stimulation increased accuracy relative to sham (Top). Stimulation did not affect reaction time (Bottom)

DCS introduces electrical stimulation artifacts in iEEG that need to be addressed before analyses can be performed. We used an ICA-based method, developed in our previous work [49], to remove the stimulation artifacts. Following artifact removal, we computed dWPLI between electrodes that were in the aWM network. As an exploratory measure, we computed magnitude-squared coherence, which is used widely in connectivity analysis of oscillatory networks. Coherence provides a complementary measure of functional connectivity as it accounts for the correlations in spectral power which is not captured by dWPLI. We restricted our analysis to the bands around the stimulation frequency for each individual participant. Additionally, we used a permutation-based approach to identify those network connections that exhibited statistically significant pairwise-differences between the conditions (in-phase stimulation vs sham stimulation, anti-phase stimulation vs sham stimulation, and in-phase stimulation vs anti-phase stimulation). This resulted in a network with sparse connections between regions within the WM network. The network connections obtained from coherence and dWPLI at the stimulation frequency for the 3 participants were pooled together for visualization in a chord diagram (Figure 4A). The nodes of the diagram represent an individual electrode while the edge between the nodes indicate the pair-wise difference in connectivity measure metric (coherence or dWPLI). Both in-phase stimulation and anti-phase stimulation resulted in more connections showing increased dWPLI than decreased, relative to sham. In contrast, the number of connections showing an increase were approximately equal to those showing a decrease when in-phase stimulation was contrasted against anti-phase stimulation. A similar trend was observed in connectivity obtained from coherence. To quantify the observed trend, we fit separate linear mixed models for pairwise difference in dWPLI and coherence with comparison as fixed factor and participant as random factor. In both cases, there was a significant effect of comparison on pairwise difference (dWPLI: F_2, 486.18_ = 26.921, p < 0.001; coherence: F_2, 533.38_ = 16.392, p < 0.001). Post hoc analysis using Tukey method revealed that the pairwise difference for in-phase vs sham and anti-phase vs sham were higher than in-phase vs anti-phase (Figure 4B, p < 0.001). The results from dWPLI and coherence suggest that contrary to our initial hypothesis, both in-phase stimulation and anti-phase stimulation increased functional connectivity relative to sham while there was no clear difference between the two stimulation conditions.

**Figure 4:**
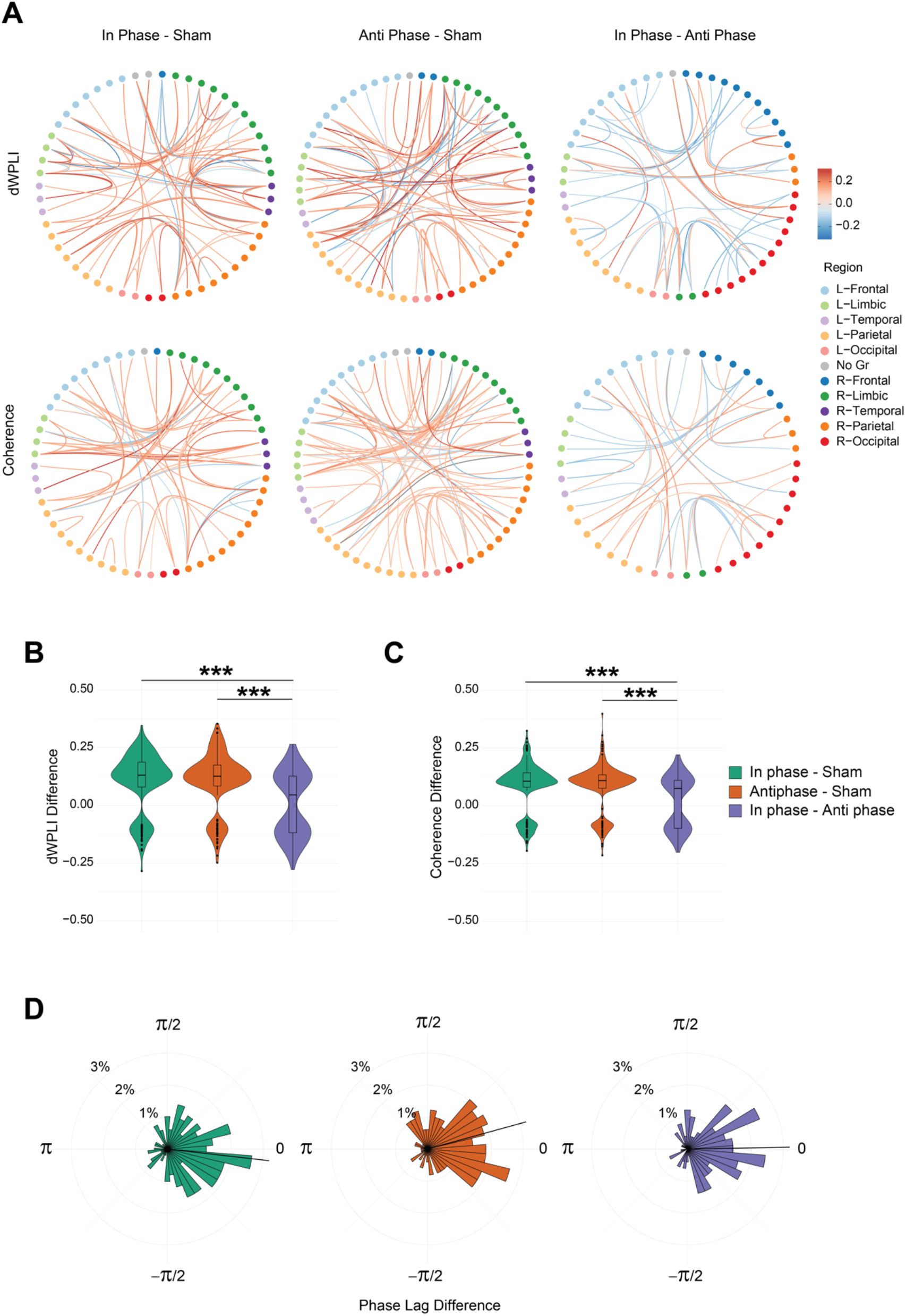
Effect of network-targeted stimulation on WM Network. A. Chord diagrams representing the pairwise differences in dWPLI and coherence in the WM network across all three participants. The nodes represent electrodes; edges represent connectivity between regions; red edges denote a relative increase in the connectivity metric and blue edges denote a relative decrease in connectivity metric between the nodes. The edges depicted here have passed a permutation based statistical significance test (p < 0.05) B. Pairwise difference of dWPLI between stimulation conditions. In-phase vs sham and anti-phase vs sham were higher than in-phase vs anti-phase differences. *** denotes statistical significance at p < 0.001 in a Tukey post-hoc test C. Pairwise difference of coherence between stimulation conditions. In-phase vs sham and anti-phase vs sham were higher than in-phase vs anti-phase differences. *** denotes statistical significance at p < 0.001 in a Tukey post-hoc test D. Circular histogram denoting the pairwise differences in phase lag across the three comparisons for the three participants. Black line denotes the mean phase lag difference for each comparison.

While these results may appear counter-intuitive, it should be noted that dWPLI is a measure of phase consistency and does not include any information regarding the actual phase difference. It is conceivable that both in-phase stimulation and anti-phase stimulation successfully engage the network due to the repeated periodic perturbation of the network and increased overall phase consistency. However, since the stimulation differed in phase lag between the targeted electrode pairs, in-phase and anti-phase may have impacted the specific phase lag between nodes in the network. To verify this, we computed phase lag at stimulation frequency between electrode pairs that exhibited significant pairwise dWPLI difference between any of the three stimulation conditions i.e., phase lag corresponding to the edges depicted in the dWPLI chord diagram in Figure 4A. Phase lag was computed from the cross-spectrum of the iEEG signal during the stimulation epoch. We pooled the data of the three participants together, as the distribution of phase lag for individual participants did not satisfy the assumptions required for the circular statistics. There was a significant effect of comparison on the phase lag differences (Figure 4C; Watson-william test F_2,488_ = 3.6523, p = 0.0266). We found that in-phase stimulation resulted in an overall decrease in phase lag relative to sham (−0.114 ± 1.310 radians; mean ± sd) while anti-phase stimulation resulted in an overall increase in phase lag relative to sham (0.269 ± 1.240 radians). There was a negligible change in phase lag when in-phase stimulation was compared to anti-phase stimulation (0.018 ± 1.165 radians). These results indicate that while in-phase stimulation and anti-phase stimulation both increased phase consistency, they modulated phase lag in opposite directions.

## Discussion

In this study, we employed a network-targeted stimulation approach to engage the WM network and test if this approach can improve WM performance. We identified oscillatory networks underlying WM from iEEG using a phase synchronization measure and stimulated functionally connected electrodes. We found in-phase stimulation improved WM performance relative to sham stimulation in all 3 participants. Interestingly, we found that both in-phase and anti-phase stimulation increased functional connectivity relative to sham. However, the effect of the two stimulation conditions on phase lag was opposite such that in-phase stimulation decreased phase lag and anti-phase stimulation increased phase lag relative to sham. The increased functional connectivity from in-phase stimulation and anti-phase stimulation may have been due to the periodic input of DCS into the WM network that aligned the phase of electrical activity between multiple regions albeit at different lags. Our results suggest that the differential effect on phase lag may have contributed to the behavioral modulation. Phase synchronization has been hypothesized to enable interareal communication by aligning periods of excitability across regions or by enabling spike timing dependent plasticity [52]. Our electrical stimulation occurred at a time scale faster than the typical timeframe for observing plasticity, suggesting that our in-phase stimulation may have aligned periods of excitability across regions that enabled enhanced communication. While in-phase stimulation improved performance, we did not observe any impairment in performance with anti-phase stimulation. In-phase stimulation may have reduced communication delay within an optimal window in which information may be effectively transmitted between regions resulting in improvement in performance. In contrast, anti-phase stimulation may have resulted in increased communication delay outside this optimal window which is inconsequential for information transmission and integration. Further studies are required to confirm this specific hypothesis.

Our results follow tACS studies that have shown behavioral effects of stimulation in WM tasks, albeit we observe improvements in accuracy while improvements in reaction times are more commonly reported. Polania et al. [34] observed a decrease in reaction time with in-phase stimulation and an increase in reaction time with anti-phase stimulation relative to sham. Violante et al. [35] observed a decrease in reaction time for in-phase stimulation relative to sham and anti-phase stimulation while there was no difference between sham and anti-phase stimulation, similar to what we observe. In addition, Violante et al. report increased BOLD signal functional connectivity increases in WM network for both in-phase and anti-phase stimulation. Although the functional connectivity from BOLD signal quantifies interactions at a slower timescale relative to what is observed in iEEG, these results support our observation that both in-phase and anti-phase stimulation resulted in increased functional connectivity.

We used a hybrid data-driven approach to restrict our analyses to putative WM networks in the three participants. The use of atlas-based priors allowed us to control the dimensionality of our variable of interest, which is the functional interactions between brain regions involved in WM; and permutation-based statistics allowed us to account for false positives. The WM network atlas we used was derived from a meta-analysis of 1091 studies that localized regions that show consistent activation across a variety of WM studies [51]. However, it must be noted that BOLD activity of regions often corresponds to iEEG activity in the high frequency broadband activity (30 – 130 Hz) [53] with lower correlations between lower frequency band activity. Therefore, it is conceivable that we may have excluded regions that exhibited task-related connectivity. Given the heterogeneity and the small sample size, we motivated this decision as a necessary trade-off for generalizability at the cost of an exhaustive naïve data-driven approach. Even so, we found task-related functional connectivity between regions in or near the aWM network in participants P1 and P2. In P1, we found theta band connectivity between superior frontal gyrus and precentral gyrus. Although superior frontal gyrus is not a part of the aWM network, our previous work has shown it may indeed play a role in WM. In P2, we found alpha band connectivity within the aWM network and stimulation resulted in the highest improvement among the three participants. In P3, even with electrodes close to the aWM network, we did not observe any significant functional interaction. This may be due to variability in the functional recruitment of brain regions for this participant. While stimulation resulted in an improvement in WM accuracy, the effect in P3 was weaker than the other two participants presumably due to the decreased recruitment of these regions by the task. However, the region of cortex activated by intracranial direct cortical stimulation extends farther than the immediate vicinity of the stimulation electrodes on the order of ~50 mm^3^ [54, 55] while the spatial extent of LFP recordings is a few millimeters [56]. Thus, stimulation may have spread into neighboring regions that are known to be canonically activated by WM task demands.

Oscillations in the theta and alpha frequency bands have been shown to support WM in many studies [57, 58] with increased oscillatory power as a marker for synchronization. Phase synchronization between brain regions in alpha and theta frequency bands have been shown to underlie many memory processes (see reviews [52, 59]) with theta band activity implicated in top-down control [34, 60, 61] and alpha band activity implicated in suppression of irrelevant information [62, 63]. Theta band synchronization has been observed between fronto-temporal and fronto-parietal regions in working memory tasks [64–67]. While fronto-parietal synchronization in alpha band has been associated with cognitive control and visuospatial attention [68], interareal synchrony has been observed to be modulated by WM load in the retention period [17, 69]. Supporting these observations, we found alpha and theta band connectivity in our participants. The variability in the frequency at which interaction was found may be due to differences in strategy [70], with theta being dominant in strategies where sequential information is encoded while alpha being dominant in strategies where competing information is suppressed. Alternately, the differences could be driven by the difference in regions between which functional connectivity is observed. We observed theta between electrodes within frontal regions while alpha was observed between electrodes in frontal and parietal regions. The studies mentioned above are constrained by the limitations of EEG, which has poor spatial resolution and is highly susceptible to volume conduction. The use of iEEG and dWPLI enabled us to address these limitations and provide a more fine-grained picture of the functional interactions.

While these results provide important insight into the role phase lag may play in coordinating working memory, the heterogeneity and the small sample size limits the interpretation to a general population. Additionally, the phase lag between stimulation sites was not taken into consideration as the initial hypothesis was based on the consistency of phase synchronization. In contrast to our approach, Kim et al. [47] stimulated hubs of a memory retrieval network at the phase lag observed between the two nodes but found that stimulation impaired performance. Our results imply that choosing a phase lag that is shorter than the observed phase lag may be beneficial. The choice of stimulation parameters was limited to in-phase stimulation and anti-phase stimulation to ensure enough trials in each condition for statistical analysis. However, this meant that we were not able to directly confirm if the effect is frequency-specific. Further studies incorporating arrhythmic stimulation as used in some TMS studies can be used to establish the frequency specificity of stimulation effects [40, 71].

While many studies quantify phase synchronization as consistency in phase differences, very few studies have focused on the phase lag between regions [34]. Given recent findings on phase-dependent information processing [72, 73] our result highlights the importance of considering phase information when studying functional interactions between brain regions. Overall, these findings advance our understanding of network-targeted stimulation for improving cognition in humans. Our results provide causal evidence that networks of brain regions are critical to cognition [74, 75] and optimal stimulation may require multi-site stimulation. This work may ultimately lead to therapeutic benefits for cognitive deficits that accompany many neurological and psychiatric disorders.

## Methods

### Participants

All experimental procedures were approved by the Institutional Review Board of University of North Carolina at Chapel Hill and informed consent was obtained from participants. Participants were recruited by invitation from patients who underwent invasive monitoring for epilepsy surgery planning. The participant clinical information is provided in Table 1. The location of electrodes in all participants were completely dictated by the clinical needs of the individual participant. See Figure S1 for the electrode coverage.

**Table 1:**
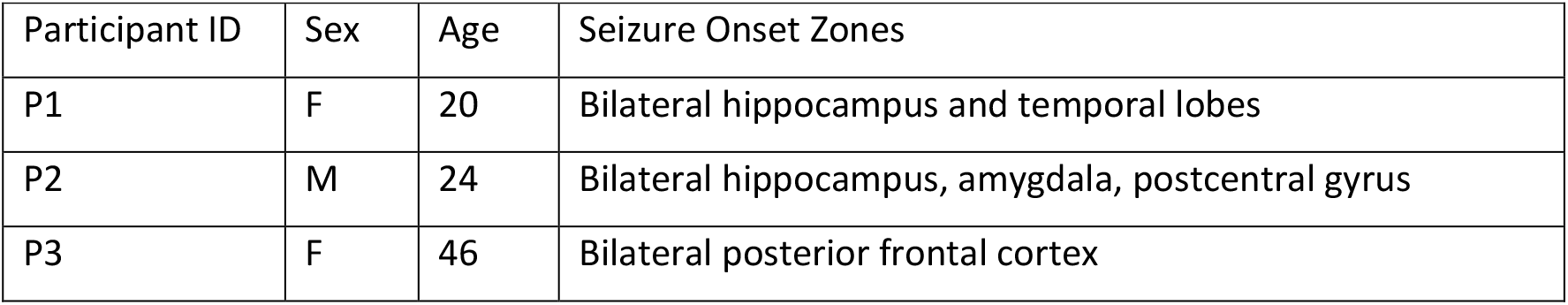
Clinical Information of Participants

| Participant ID | Sex | Age | Seizure Onset Zones |
| --- | --- | --- | --- |
| P1 | F | 20 | Bilateral hippocampus and temporal lobes |
| P2 | M | 24 | Bilateral hippocampus, amygdala, postcentral gyrus |
| P3 | F | 46 | Bilateral posterior frontal cortex |

### Working Memory Task

Participants performed a Sternberg working memory task that has been previously used in ECoG studies [11, 49, 76]. The Sternberg task allows a separation of different cognitive processes involved in working memory into different epochs: encoding, maintenance, and retrieval (Figure 1 B). Each trial began with a fixation cross presented for 1000 ms. In the encoding epoch, participants were presented with a sequence of letters from the English alphabet one letter at a time. Each letter was presented for 500 ms. Following the encoding epoch, a blank screen was presented for 2000 ms which served as the maintenance epoch. Next, a single letter (probe) was presented on the screen for 3000 ms. The participants were instructed to indicate if the probe was present in the encoding epoch or not using custom joysticks that interfaced with the task administration laptop through a USB response box (Black Box Toolkit, Sheffield, UK). P3 was not able to use the joysticks due to history of stroke affecting motor function in their right hand and responded using the keyboard of the laptop with their left hand only. The task was programmed in Matlab using Psychtoolbox [77].

Participants completed the task in two sessions – a baseline session and a stimulation session. In the baseline session, the task consisted of memory arrays of two different lengths (WM load). In the stimulation session, the WM load was fixed to maximize the number of trials in each stimulation condition. The experimental parameters used for the participants are listed in Table 2.

**Table 2:** Experimental Parameters for the Participants

| Participant ID | Baseline |  | Stimulation |  | Stimulation Electrode Pairs Location | Stimulation Frequency |
| --- | --- | --- | --- | --- | --- | --- |
|  | WM Load | No. Trials/Load | WM Load | No. Trials/Condition |  |  |
| P1 | 3,5 | 30 | 5 | 30 | Left anterior superior frontal gyrus, Left inferior precentral sulcus | 4 Hz |
| P2 | 5,7 | 40 | 7 | 30 in-phase, 36 anti-phase, 34 Sham | Left inferior frontal junction, left superior parietal lobule | 10 Hz |
| P3 | 5,7 | 40 | 5 | 40 | Right anterior middle frontal gyrus, right superior intraparietal sulcus | 10 Hz |

### ECoG Data Acquisition and Direct Cortical Stimulation

ECoG data were recorded using a 128-channel EEG system (NetAmps 410, Electrical Geodesics Inc, Eugene, Oregon, United States) at 1000 Hz sampling rate. Stimulation was delivered using Cerestim M96 cortical stimulator (Blackrock Microsystems, Salt Lake City, Utah, United States). Stimulation consisted of a train of biphasic pulses 2 mA in amplitude, 200 μs in duration per phase of the biphasic pulse with a 55 μs interval between the positive going and negative going phase. The inter-pulse-interval was adjusted according to the stimulation frequency. Stimulation was applied between two pairs of electrodes identified from functional connectivity analysis as described in the next subsection. The timing of pulses between the two pairs was in-phase, i.e., stimulation was applied simultaneously between the two electrode pairs (Figure 1C). We hypothesized that in-phase stimulation would improve WM performance. The active control was anti-phase stimulation, i.e., stimulation between the first pair and second pair was offset by half the inter-pulse-interval (Figure 1C). Both in-phase and anti-phase stimulation was time-locked to the start of the encoding epoch. Stimulation was triggered using Matlab wrapper functions provided by the manufacturer of the cortical stimulator. In addition, a control condition where no stimulation was applied (sham) was also included to account for any non-specific effects of stimulation.

### Data Analysis

All data analysis was performed using custom written Matlab scripts utilizing functions from the EEGLAB [78] and Fieldtrip toolboxes [79]. Electrodes over seizure focus were excluded from analysis.

### Selection of Stimulation Electrodes

ECoG data collected during the baseline session was used to determine functionally connected electrodes. The continuous data was band-pass filtered between 1 and 50 Hz using an FIR filter and re-referenced to the average of all intracranial electrodes using functions from EEGLAB toolbox. The data was then segmented into trials containing the different epochs. Functional connectivity was determined using debiased weighted phase lag index square (dWPLI) implemented in Fieldtrip toolbox. The measure is a composite of phase lag index, which captures consistency in phase lag between two time oscillatory signals [80], and the imaginary part of coherence which ignores zero phase lag interactions [81]. DWPLI has been shown to provide a better estimate of phase-synchronization in the presence of volume conduction and the debiased estimate has higher statistical power [50]. DWPLI was computed for the fixation, encoding and retention epochs separately. The strength of functional connectivity was strongest between neighboring electrodes followed by electrodes within the same anatomical region, i.e., frontal cortex or parietal cortex. Since we were interested in modulating long-range functional connectivity, we ignored electrode pairs that were neighbors. In addition, connections that were present in the fixation epoch and between electrodes over seizure foci were ignored as the former may reflect preparatory attentional components of network activity and the latter may reflect pathological connectivity.

### Removal of electrical stimulation artifacts

Electrical stimulation artifacts were removed using an independent component analysis (ICA) based approach as demonstrated in our previous work [49]. Artifacts appear as stereotypical waveforms in iEEG signals. Blind signal separation using ICA separates the iEEG signal into components that contain only artifact waveforms and other components that contain the rest of the signal. The components containing artifacts were then rejected and the remaining components were used to reconstruct the artifact free signal. We used the infomax algorithm [82] available as a part of EEGLab toolbox for computing independent components. Following artifact suppression, the signals were re-referenced to the average of all signals.

### Estimation of functional connectivity

DWPLI was computed for the stimulation session (epoched by stimulation condition) in the same manner as the baseline session (epoched by WM load) using functions from EEGlab and Fieldtrip toolboxes. In addition, coherence was also computed for the stimulation session. Adjacency matrices were derived from a 3 Hz band centered on the frequency of interest. Phase lag was derived from the mean cross-spectrum across trials in a 2 Hz band centered on the frequency of stimulation. For pairwise comparisons between stimulation conditions, the difference in adjacency matrices were computed. Statistical significance was computed using a permutation-based approach. Trial labels were shuffled 1000 times, and adjacency matrices were computed for each condition. Pairwise differences were computed as above to generate a null distribution. Any pairwise difference in the non-shuffled adjacency matrices that were greater (or lesser) than 95% of the null distribution differences were deemed statistically significant. Chord diagrams were plotted using ggraph and igraph packages written in R.

### Identification of electrode locations

3D Slicer [83] was used to analyze and extract electrode locations from CT images obtained after implantation of subdural electrodes (post-OP CT). Electrode locations were determined manually using the post-OP CT image by placing fiducials in areas of high activation in the CT. The post-OP CT was co-registered to pre-OP MRI in Slicer. The anatomical locations of the electrodes were determined by co-registering the pre-OP MRI Image to the MNI Atlas [84], recomputing electrode locations in the MNI space, transforming these locations to Talairach space, and using the Talairach Client [85] to obtain the label of the region nearest to the coordinate representing electrode location.

### Determining atlas-based WM network

We used a meta-analysis-based approach to identify regions activated by a variety of WM tasks. Using the Neurosynth database, we acquired the association test map for ‘Working Memory’ that was derived from 1091 studies. The map consisted of z-scores, corrected with false discovery rate (FDR) at an alpha value of 0.01, from a two-way ANOVA testing for the presence of a non-zero association between the term ‘Working Memory’ and voxel activation [51]. We defined 8 mm regions of interest (ROIs) around each electrode in the Montreal Neurological Institute (MNI) space using custom written scripts and the MarsBaR toolbox in SPM12 [86, 87]. Next, we determined the z-scores from the association map within these ROIs (electrodes) and computed the mean z-score for each electrode. Any electrode that had a mean z-score greater than 0 was defined to be part within the aWM network and was included in analysis.

### Statistical Analysis

All statistical analyses were performed using custom-written scripts in R. Linear mixed models were fitted using *‘lmertes’* package [88]. The package uses a Sattherwaite approximation for degrees of freedom for ANOVA. Post-hoc analyses consisted of pairwise comparisons with Tukey adjustments and were computed using *‘emmeans’* package. Circular statistics were computed using *‘circular*’ package [89].

## Acknowledgments

The authors thank the members of the Frohlich Lab for their valuable input. The authors also thank the EEG technicians at the UNC Epilepsy monitoring unit for their generous help with the iEEG recordings.

## Funding

Research reported in this publication was supported in part by the National Institute of Mental Health of the National Institutes of Health under Award Numbers R01MH101547 and R21MH105557, National Institute of Neurological Disorders and Stroke of the National Institutes of Health under award number R21NS094988-01A1. The content is solely the responsibility of the authors and does not necessarily represent the official views of the National Institutes of Health.

## Supplementary Information

**Figure S1:**
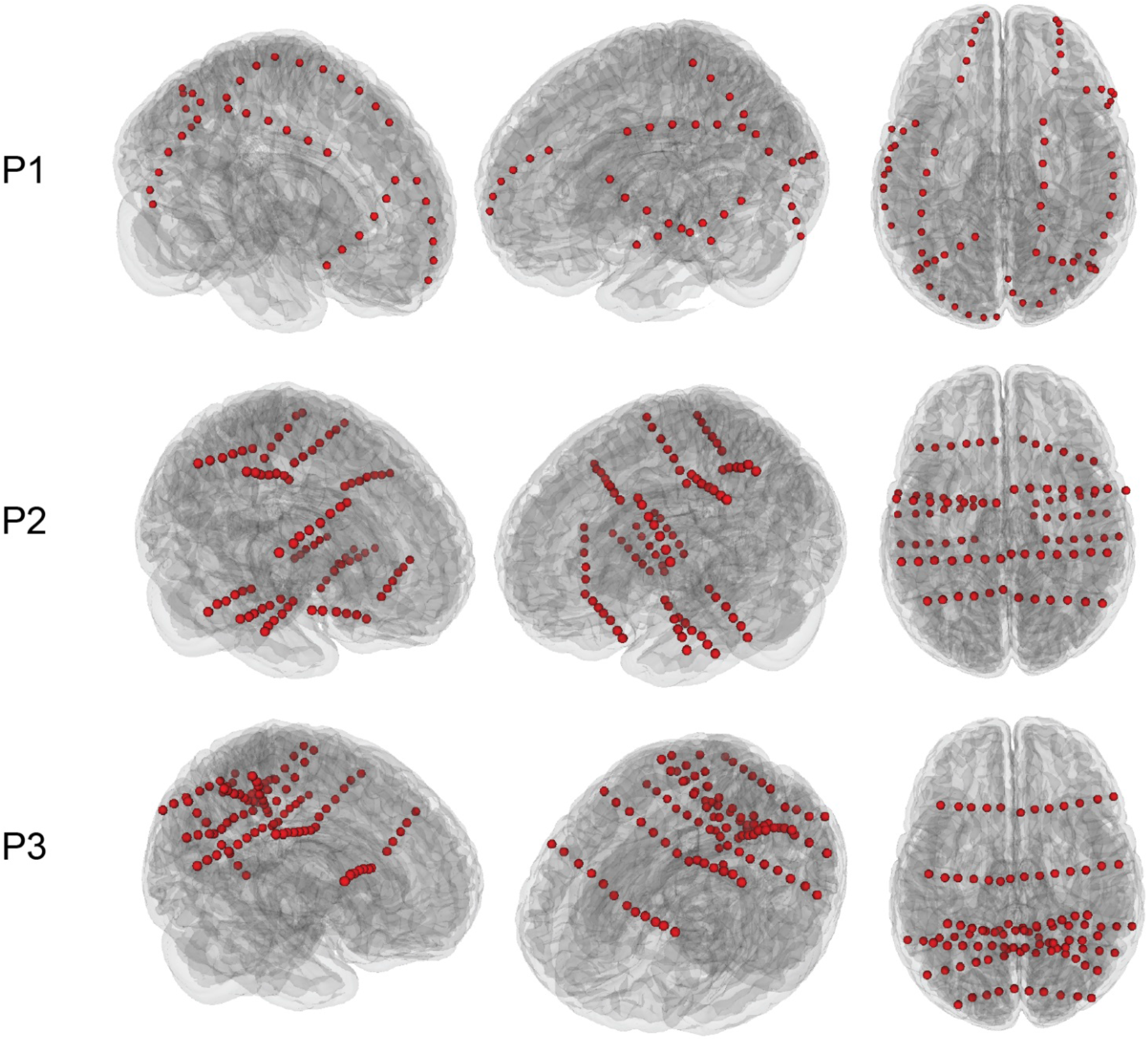
Locations of implanted electrodes in the three participants. Red circles denote the electrodes implanted in the three participants. Subdural strip electrodes were used in P1 while stereotactic EEG electrodes were used in P2 and P3

**Figure S2:**
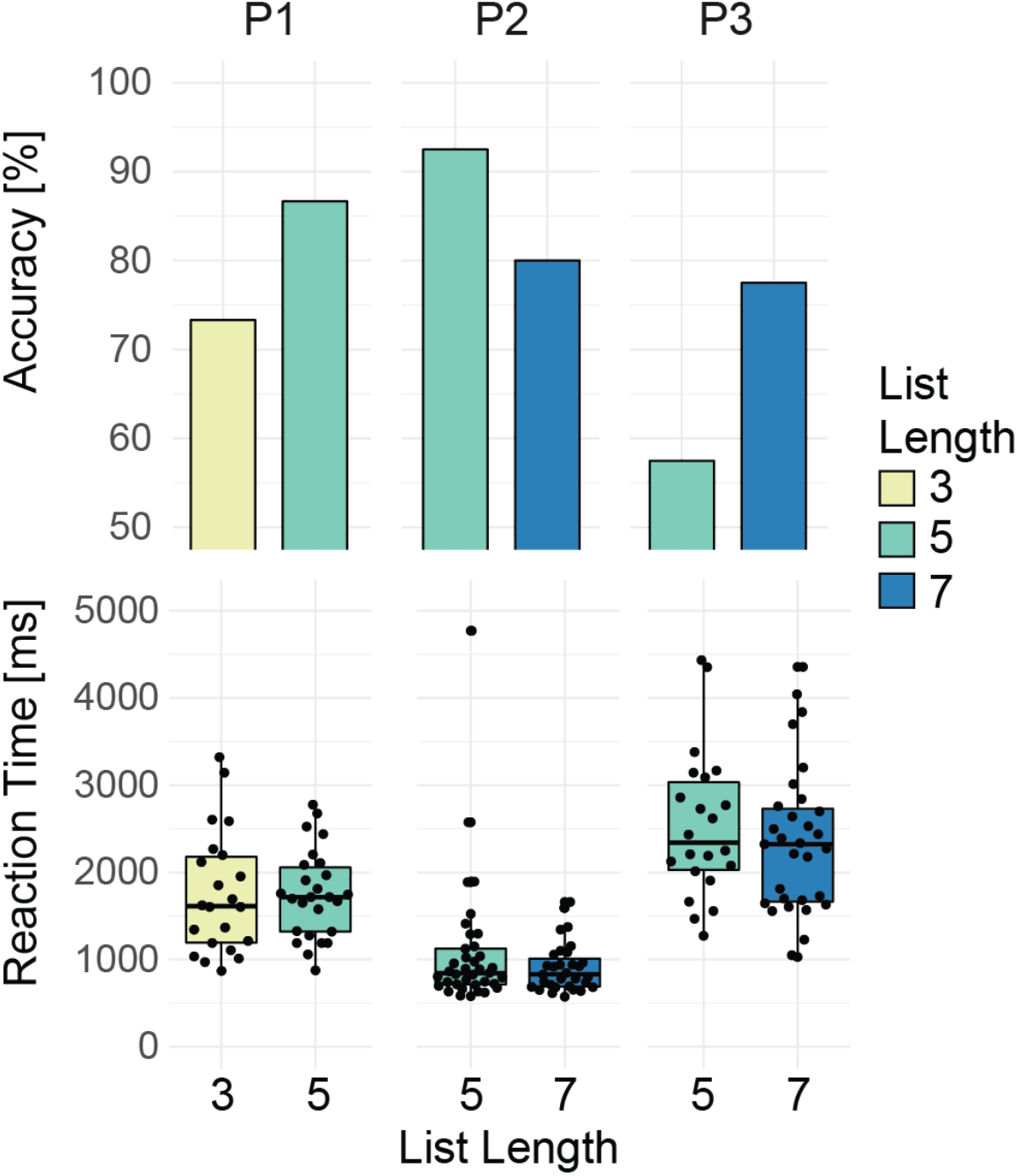
WM performance in baseline session. Accuracy and reaction time (in correct trials) were not statistically significant with increased list length.

## Notes

Conflict of interest: FF is the lead inventor of IP filed on the topics of noninvasive brain stimulation by UNC. FF is the founder, CSO and majority owner of Pulvinar Neuro LLC, which played no role in this research The other authors declare no competing interests

